# Morphine Suppresses Peripheral Responses and Transforms Brain Myeloid Gene Expression to Favor Neuropathogenesis in SIV Infection

**DOI:** 10.1101/2022.07.25.501436

**Authors:** Howard S. Fox, Meng Niu, Brenda M. Morsey, Benjamin G. Lamberty, Katy M. Emanuel, Palsamy Periyasamy, Shannon Callen, Arpan Acharya, Gregory Kubik, James Eudy, Chittibabu Guda, Shetty Ravi Dyavar, Courtney V. Fletcher, Siddappa N. Byrareddy, Shilpa Buch

## Abstract

The twin pandemics of opioid abuse and HIV infection can have devastating effects on physiological systems, including on the brain. Our previous work found that morphine increased the viral reservoir in the brains of treated SIV-infected macaques. In this study, we investigated the interaction of morphine and SIV to identify novel host-specific targets using a multimodal approach. We probed systemic parameters and performed single-cell examination of the targets for infection in the brain, microglia and macrophages. Morphine treatment created an immunosuppressive environment, blunting initial responses to infection, which persisted during antiretroviral treatment. Antiretroviral drug concentrations and penetration into the cerebrospinal fluid and brain were unchanged by morphine treatment. Interestingly, the transcriptional signature of both microglia and brain macrophages was transformed to one of a neurodegenerative phenotype. Notably, the expression of osteopontin, a pleiotropic cytokine, was significantly elevated in microglia. This was especially notable in the white matter, which is also dually affected by HIV and opioids. Increased osteopontin expression was linked to numerous HIV neuropathogenic mechanisms, including those that can maintain a viral reservoir. The opioid morphine is detrimental to SIV/HIV infection, especially in the brain.

## INTRODUCTION

The clinical outlook for people with HIV (PWH) has improved dramatically as increasingly effective combination antiretroviral therapy (cART) has become available (1–4). Since untreated HIV infection leads to the acquired immunodeficiency syndrome (AIDS), a focus on the immune abnormalities has been prominent. However, the central nervous system could also be adversely affected. Before the onset of successful treatment options, 8-15% of PWH in the United States (US) developed a severe cognitive disorder, HIV-associated dementia (HAD) (5, 6). In the post-ART era HAD has been reduced to 1-2% in the US and 5% globally (7, 8). Paradoxically, increased survival rates resulting from therapy have also resulted in an undesirable increase in the prevalence of minor neurocognitive impairments, collectively referred to as HIV-associated neurocognitive disorders (HAND), also known as neuroHIV (7, 8). Unfortunately, cART does not completely shield the brain from the effects of HIV-1 infection.

HIV, as well as the analogous nonhuman primate (NHP) virus SIV, enters the brain early after infection (9–13), and can set up a lifelong presence in the CNS (14, 15). Many widely used antiretroviral drugs poorly penetrate the CNS, perhaps facilitating pathogenic effects of infection and the maintenance of a viral reservoir (16). HIV infection remains a significant problem: at the end of 2018, there were over one million people diagnosed with HIV infection in the US and an estimated over 160,000 people whose infections had not been diagnosed (17).

Over two million people in the US have opioid use disorder (OUD), a significant risk for HIV infection and transmission; in addition, PWH are frequently prescribed opioids for pain management (18). Opioid abuse and HIV have been described as dual linked health crises (19), and drug abuse could likely be one of the leading causes of HIV transmission in the United States. Injection drug use was found to be responsible, solely or in conjunction with other causes, for 11% of HIV infections in 2020 (20), and other cases can arise through risky sexual behavior due to drug use (21). While HIV infection has transformed from a death sentence to a manageable chronic condition, the comorbid condition of drug use, such as opioid abuse, in infected individuals is on the rise, and may lead in turn to increased neurologic and cognitive deficits in PWH (22). The CNS may be uniquely susceptible to the combined effects of opioid abuse and HIV. Among the various abused drugs, the use of opioids is highly prevalent in PWH, both from illicit drug use and the management of HIV-associated neuropathic pain. It is estimated that one-quarter to one-half of PWH are prescribed opioids, and PWH are prescribed higher doses than those without HIV (23). Intriguingly, brain regions that are targets for morphine (expressing high numbers of μ-opioid receptors), are also the regions with increased viral loads and predilection for HIV infection (24–26). Thus, the interplay of HIV and opioids raises concerns regarding the effect of opioids on HIV’s impact on the brain. Identifying CNS molecular aberrations underlying HIV and opioid abuse thus remains an unmet need in the field.

Regarding neuroHIV, the effects of substance abuse can vary (27, 28). Studies from both before and after the introduction of treatment have reported increased neuropathogenesis associated with opioid use (29, 30). Still, studies on opioid users as well as results on the effects of opioids on HIV studied through *in vivo* and *in vitro* systems have yielded varied results on whether opioids increase neuroHIV or factors attributed to HIV neuropathogenesis; these have been thoroughly reviewed recently (18, 22, 31). The different findings could be attributed to the pleiotropic effects of opioids, such as in the balance of opioid-enhanced infection by the virus versus opioid-induced dampening of inflammatory lesions. In addition, the conflicting findings in humans could be attributed to high variability in drug usage patterns, use of different opioids, multidrug use, differences in nutritional and socio-economic status, variable access to medical care, and potential inaccuracy in drug use data collection due to overdependence on self-reporting. In the current era issues such as access for treatment in opioid users due to economic and social issues, difficulties in treatment compliance due to drug use and its consequences, such as incarceration, and comorbid conditions due to drug use and additional complicating issues.

NHP infection by SIV remains the best model for the study of HIV pathogenesis and treatment, including neuropathogenesis (32–37). NHPs are also an excellent animal model for studying the effects of drugs of abuse and their interactions with SIV infection, especially for effects on the brain (38–42). Such animal experiments allow better controlled conditions than observational studies in humans. However, studies on the effects of opioids during experimental infection of NHP by SIV in the absence of treatment have yielded inconsistent findings (39, 41, 43–46).

In SIV-infected NHP, as in people, the viral targets in the brain are primarily brain macrophages and microglia (47, 48). These key cells support productive HIV and SIV replication in the CNS and can serve as virus reservoirs in the brain (48–52). Infection of macrophages and microglia and the subsequent inflammation associated with infection can alter their function and phenotype, affecting other glia as well as neurons. In parallel, opioids have also been shown to impact macrophage and microglial functioning and phenotype (53–56).

We recently reported that in monkeys chronically administered the prototypic opioid morphine, followed by SIV infection and subsequent suppression of viremia below the limit of detection by cART, the size of the viral reservoir in the brain was increased relative to that in animals given saline instead of morphine (57). In the current study, we investigated longitudinal changes in response to SIV infection of rhesus monkeys in the presence of morphine compared to animals administered saline before and after cART treatment. Using single-cell and single-nucleus RNA sequencing (scRNA-seq and snRNA-seq), we examined the effect of morphine on brain-resident macrophages and microglia in these SIV-infected, cART-suppressed animals to better understand the factors responsible for the morphine-induced increased CNS reservoir and the role of opioids in neuropathogenesis.

## RESULTS

### Acute cellular response to SIV is suppressed by morphine

In our recent study, we infected morphine-dependent (n=6) and saline-administered control (n=5) rhesus monkeys with SIV, followed by treatment with cART to suppress viremia, modeling the acquisition of HIV infection in those who use opioids (57). As reported previously, no differences in plasma viral load were found due to morphine, but at the 2-week post-infection time point, cerebrospinal fluid (CSF) viral load was lower in the morphine group. After 30 weeks (or more) of treatment with antiretrovirals, resulting in viral suppression in plasma and CSF, analysis of the brain viral reservoir revealed that it was higher in CD11b+ microglia/ macrophages in morphine-dependent animals (57). Numerous factors could drive this effect. To address these in the context of morphine, we first examined two general health measures and several aspects of blood cell composition in the two groups throughout infection (Figure 1). Morphine-treated animals had a significantly lower weight gain than the saline-group, consistent with the known effects of morphine on appetite. Albumin, a general marker of nutrition, remained unchanged.

**Figure 1.**
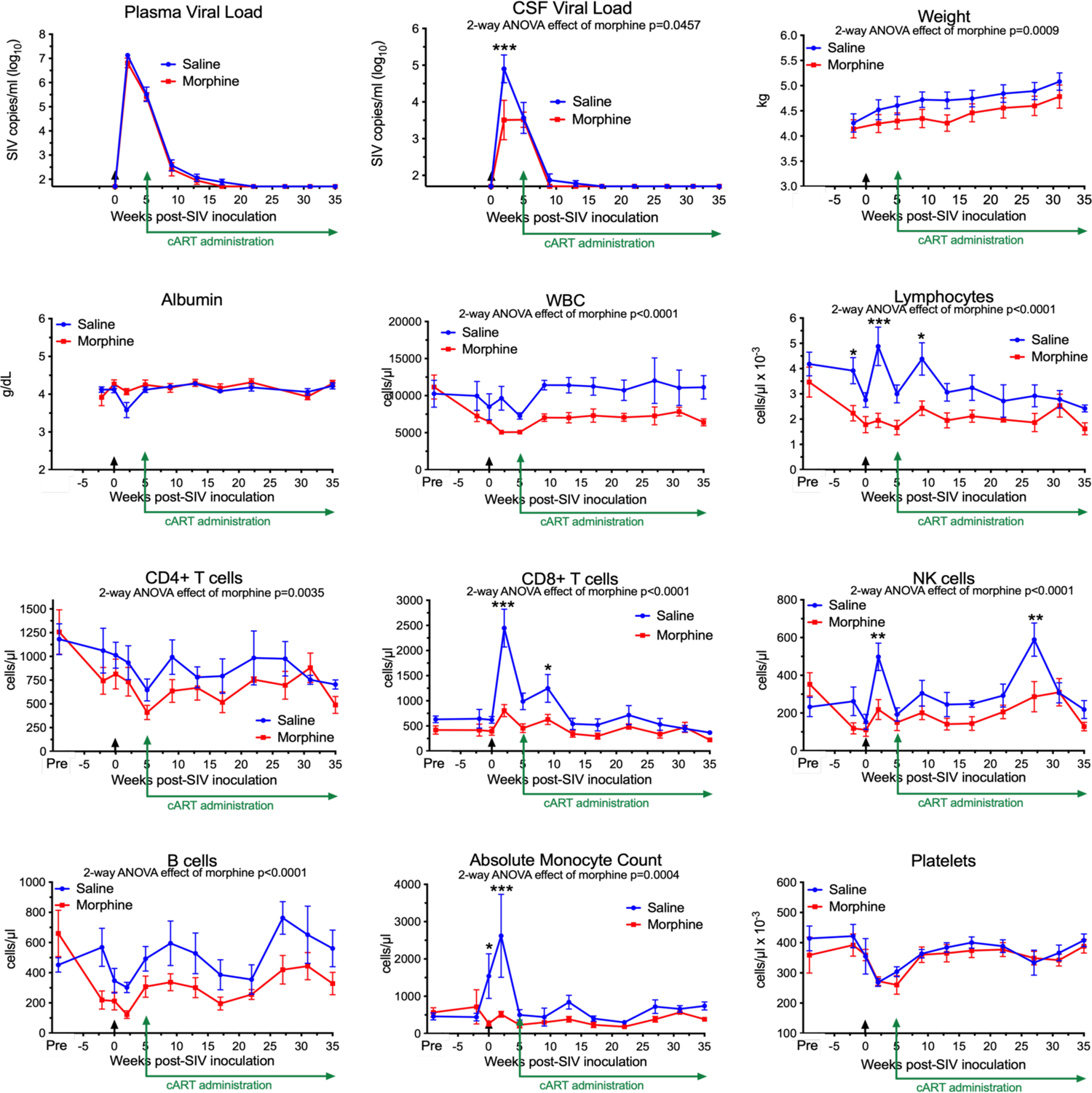
Longitudinal Assessments. Viral loads, general health, and blood cell composition in the saline (blue) and morphine (red) groups. The points represent the mean and the error bars the standard error of the mean. Viral inoculation is denoted by black arrow at week 0; cART was begun at week 5 and continued throughout the rest of the experiment. Biofluid viral loads were previously reported (57). The weight and albumin data begin at week −2, when the morphine group animals were on stable maintenance doses of morphine. The blood cell composition includes a pre-morphine value (and the equivalent time in the saline group). Statistical testing was done by 2-way ANOVA and the p-value indicated above for significant differences between the groups. When the ANOVA was significant, testing on the individual time points was performed using Sidak’s multiple comparison test. * = <0.05, ** = < 0.01, *** = < 0.001.

Striking findings were found in blood cells. The total number of white blood cells and the lymphocyte subset were significantly lower in the morphine group. All lymphocyte subsets examined: CD4+ T cells, CD8+ T cells, NK cells and B cells were lower in the morphine group. Strikingly the increase in CD8+ T cells and NK cells at the acute infection time point (2 weeks p.i.) in the saline group was significantly reduced in the morphine group. Monocytes showed the same pattern, lower in the morphine group, and the increase found in the saline group at 2 weeks p.i. largely ablated. Platelets, while dropping in the acute and post-acute infection stage, with recovery after cART administration, did not differ between the two groups. Thus the acute cellular immune reaction to infection, in terms of increases in lymphoid and myeloid cells, was significantly suppressed in response to morphine.

### Chronic morphine creates an immunosuppressive environment

Next, we addressed the expression of several blood biomarkers of various physiological processes, including inflammation. Plasma from the six morphine-administered animals was assessed pre- and post-morphine ramp-up and maintenance administration (2-week ramp-up period and 7-week maintenance), before SIV infection. Interestingly, several molecules were altered by morphine (Figure 2, left graft for each analyte). Adiponectin, a hormone made by fat cells, functions to sensitize cells to insulin as well as anti-inflammatory effects (58), was significantly increased by morphine. In keeping with this finding on an anti-inflammatory molecule, a number of inflammatory markers were decreased following morphine treatment: interleukin 16 (IL-16), a proinflammatory cytokine that is a chemoattractant for a number of different classes of immune cells (59); matrix metalloprotein 2 (MMP-2), one of a family of MMPs that serve to degrade the extracellular matrix to enable cellular inflammation (60); tumor necrosis factor receptor 2 (TNFR2), one of the two TNF receptors, and its soluble form used as a biomarker for TNF activity in inflammation (61); and vascular cell adhesion molecule 1 (VCAM-1), an endothelial adhesion molecule for immune cells, and its soluble form used a biomarker for inflammation and endothelial activation (62).

**Figure 2.**
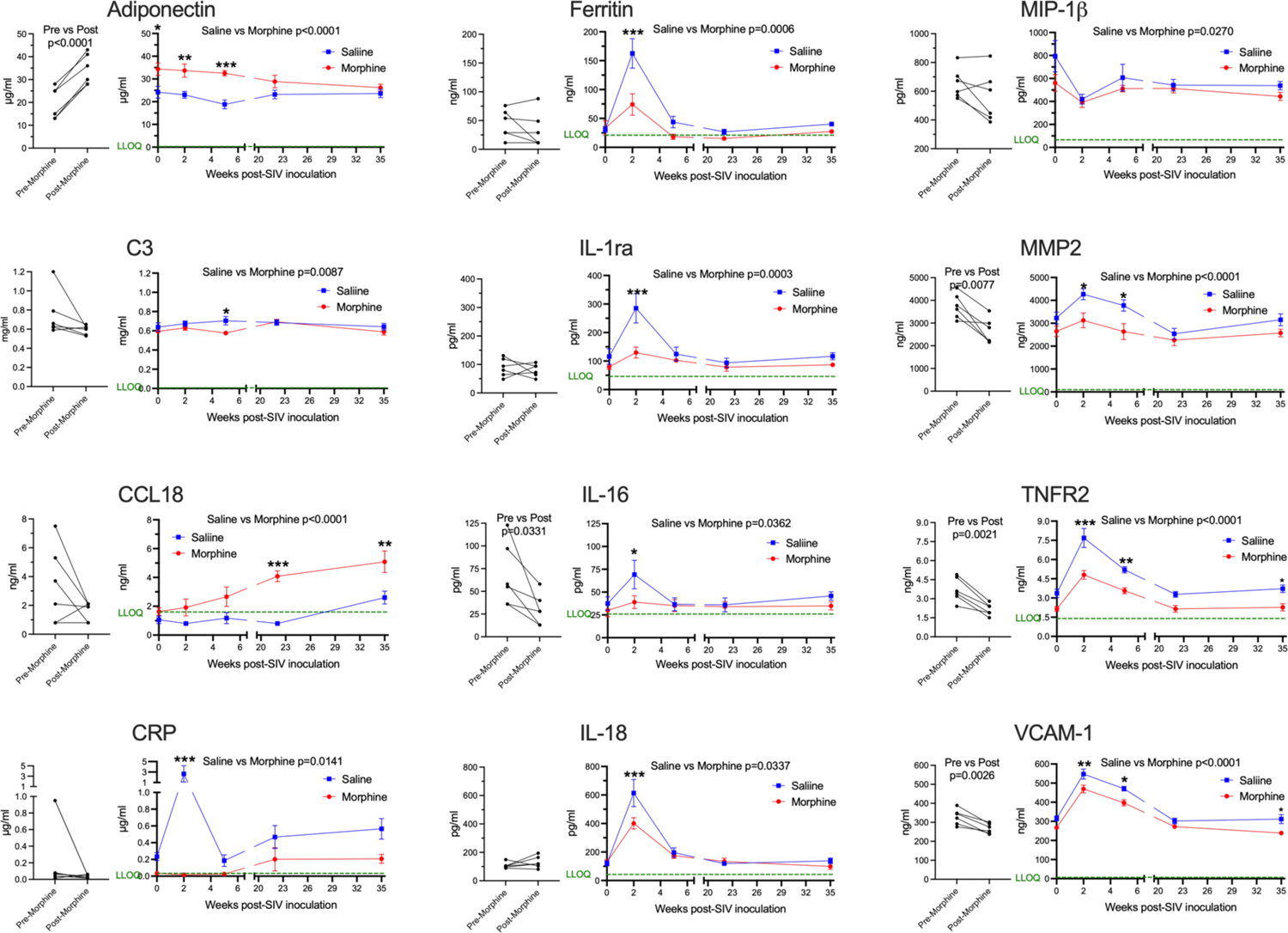
Plasma biomarkers. A panel of plasma proteins was assessed before and after morphine administration in the morphine group, and then throughout infection in both the morphine (red) and saline (blue) groups. The left panel for each analyte contains the pre- and post-morphine (before SIV inoculation); statistical testing was performed by paired t-test. The right panels show values starting on the day of inoculation (week 0), and then week 2 (acute), week 5 (immediately before cART administration began), and then with viremia suppressed by cART, week 22 and week 35. The points represent the mean and the error bars the standard error of the mean. Statistical testing was done by 2-way ANOVA and reported for significant differences between the groups. When the ANOVA was significant, testing on the individual time points were performed using Sidak’s multiple comparison test. * = <0.05, ** = < 0.01, *** = < 0.001. LLOQ = lower limit of quantitation, shown as a green line. One-half the LLOQ value was used for nonquantifiable values <LLOQ.

Following this baseline period of morphine administration, both the morphine and saline groups of animals were inoculated with SIV. In addition to the pre-inoculation time point (week 0), biomarkers were examined during the acute phase of infection at the peak of viremia (2 weeks post-infection, p.i.), when the acute viremia was lowered by the host immune response (5 weeks p.i.), at two points following virus suppression by cART (22 and 35 weeks p.i.) (Figure 2, right graph for each analyte). For the longitudinal measurements, each of the above analytes was significantly different, with adiponectin increased, and MMP-2, TNFR2, and VCAM-1 decreased. Strikingly, several additional molecules were also altered over the time course and showed specific significant differences during the acute 2-week p.i. time point. C-reactive protein (CRP), a general marker for inflammation (63), increased dramatically in saline-administered animals, whereas the levels did not change over baseline in the morphine-administered animals. Ferritin, an iron-sequestering protein, is known to be elevated in inflammation due to cellular damage (64), and increased significantly in saline compared with morphine-administered animals. The interleukin 1 receptor antagonist (IL-1ra), produced in response to inflammation to help control proinflammatory IL-1 effects (65), was also elevated to a greater extent in the saline group. Interleukin 18 (IL-18), a pleiotropic proinflammatory cytokine in the IL-1 family (65), was also elevated acutely in the morphine group.

The expression of several additional molecules was found to be altered in the morphine-treated group compared with the saline-treated group over the course of infection. Complement factor 3 (C3), which is elevated in inflammation (66), showed a slight decrease in the morphine group, reaching significance at week 5 p.i. Macrophage inflammatory protein-1 beta (MIP-1β), a pro-inflammatory chemokine for a number of lymphocyte subsets and inhibitor of HIV and SIV infection through CCR5 (67), also showed slightly decreased levels in the morphine-treated group. In contrast, C-C motif chemokine ligand 18 (CCL18) was increased over the time course of infection by morphine, particularly in the chronic virus-suppressed time points. CCL18 has been found to have both immune activating and immune suppressing functions (68). A list of the analytes measured, and their level of expression is given in Supplemental Table 1.

In summary morphine generally suppressed inflammatory markers before and after infection. These effects were magnified at the acute 2-week p.i. time point, a time point when the immune system is highly activated in response to the infection and high level of viremia. This decreased inflammatory reaction is consistent with the decreased cellular response above in the morphine group.

### CNS and peripheral antiretroviral distribution and concentration are unchanged by morphine

While the morphine immunosuppression could help explain the increased SIV myeloid reservoir present in the brain and untoward effects of opioids on HIV pathogenesis and neuroHIV, it is likely that modulation of antiretroviral (ARV) levels in biofluids and tissues by morphine can affect the reservoir size. We thus sought to measure the concentrations of ARVs in the plasma and CSF as well as tissues. At necropsy, animals were given their daily subcutaneous injection of cART, followed 90 minutes later by bleeding and CSF sampling, with subsequent tissue harvest 20-30 minutes later. No difference was found between the ARV drug concentrations in the saline or morphine groups in any of the measured parameters (Figure 3A). Some differences, however, were found among the different brain regions (Figure 3B). For example, dolutegravir (DTG) was lower in the white matter than the putamen or pons. The active triphosphate of tenofovir (TFV-diphosphate, TFV-DP) was lower in the frontal cortex than the hippocampus, putamen, pons and white matter, and the putamen lower than pons and white matter. The active triphosphate of emtricitabine (FTC-TP) was lower in the frontal cortex and putamen than in the pons. When significant differences were found, the pons had higher levels for all three ARVs within the brain. Still, the levels in the brain were consistently lower than those in the rest of the body.

**Figure 3.**
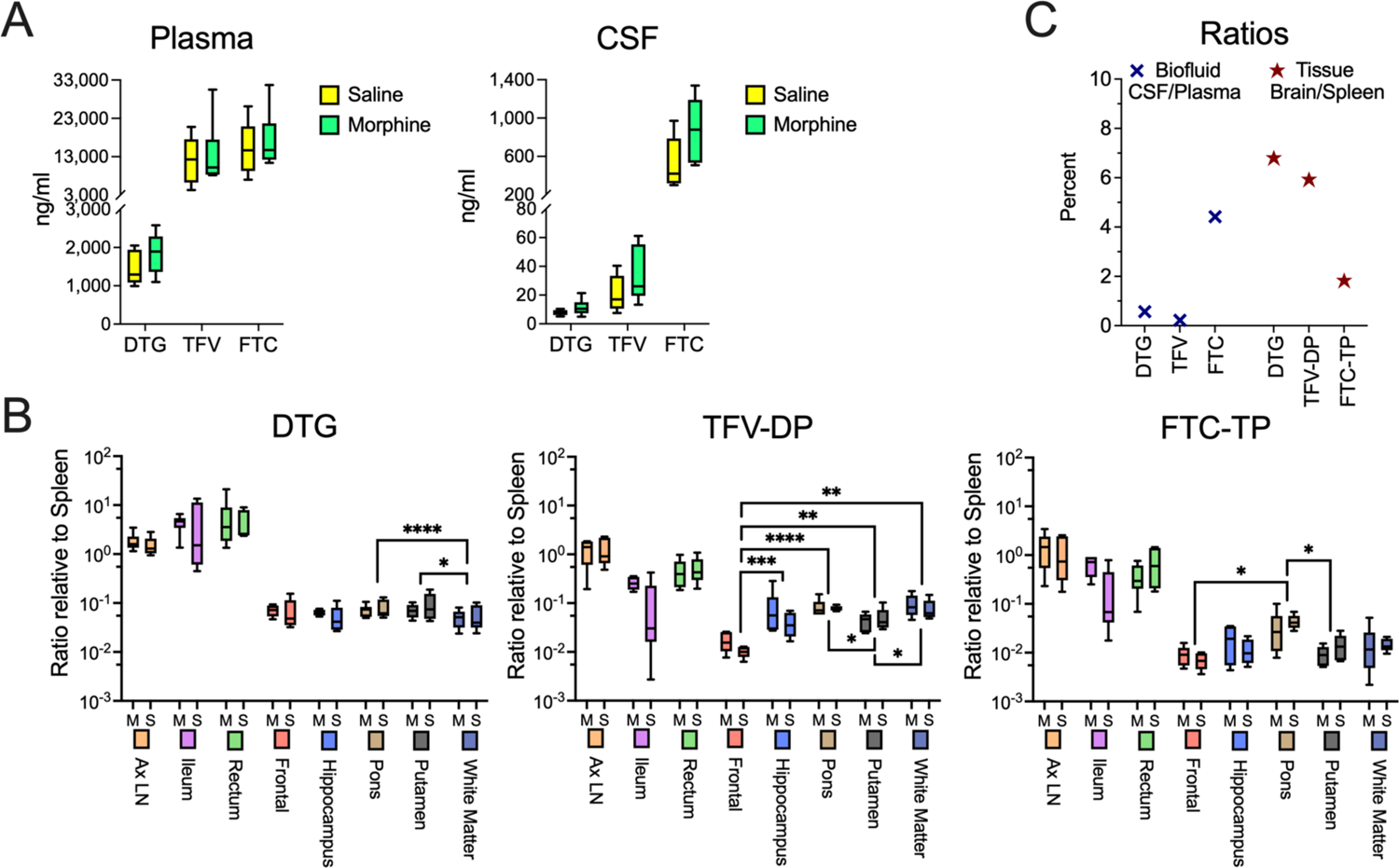
Antiretroviral drug measurements in biofluids and tissues at necropsy. **A)** Concentrations of dolutegravir (DTG), tenofovir (TFV) and emtricitabine (FTC) in the plasma and CSF, 90 minutes after subcutaneous administration of the drugs. **B)** Tissue concentrations (with brain regions indicated) of DTG and the active diphosphate (DP) metabolite of TFV (TFV-DP) and triphosphate (TP) metabolite of FTC (FTC-TP). Amounts of these entities were normalized to the amounts found in the spleen of each animal. Statistical testing was done by 2-way ANOVA for the brain region only. When the ANOVA was significant, testing on the individual regions was performed using Sidak’s multiple comparison test. * = <0.05, ** = < 0.01, *** = < 0.001. **C)** Ratio of the concentrations found in CSF relative to spleen (left) and of the average of the 5 brain regions relative to spleen (right).

Interestingly, examining CSF concentrations, FTC had the highest penetration from plasma to CSF, whereas for the active drug, when comparing spleen to brain concentrations, FTC-TP was the lowest of the three drugs (Figure 3C), consistent with our findings in rodents (69, 70). Although in the biofluids the CSF drug concentrations are much lower than those in plasma, the CSF drug concentrations exceeded the protein-free (as CSF has greatly reduced protein compared to plasma) 50% inhibitory concentration (IC_50_, as measured for HIV) for TFV (1.98-fold greater), FTC (8.59-fold greater) and the IC_90_ for DTG (13.3-fold greater). However the intracellular tissue concentrations of the active drugs were below the IC_50_/IC_90_ in the brain for the drugs, with higher values, many exceeding the IC_50_/IC_90_, in the lymphoid organs and gut (Supplemental Figure 1). Overall, there is no evidence that morphine-related differences in ARV brain concentrations affect the CNS viral reservoir or aspects of CNS function.

### Chronic morphine does not change the pattern of CNS myeloid cell subsets

Since SIV and HIV target the myeloid cells in the brain, and such cells can serve as a viral reservoir during treatment as well as drive neuropathogenesis, we next characterized the myeloid cells (microglia and macrophages) purified from the brain of four animals in each group by single-cell RNA sequencing (scRNA-seq). On average we examined ∼5,700 such cells from each animal. No expression of SIV transcripts was identified in the cells. Graph-based clustering revealed the cells could be divided into six clusters (Figure 4A). Morphine did not alter the proportion of cells in the clusters, as the distribution of cells from each condition was similar between the clusters (Figure 4B). Using a panel of myeloid gene markers, based on gene expression profiles, clusters 1-4 and 6 most closely resemble microglia, with higher C-X3-C motif chemokine receptor 1 (CX3CR1), G protein-coupled receptor 34 (GPR34), and purinergic receptor P2Y12 (P2RY12). On the other hand, cluster 5 consists of macrophages, with higher levels of CD74 (the major histocompatibility class II invariant chain), CD163, major histocompatibility complex, class II, DR alpha (MAMU-DRA), as well as S100 calcium binding protein A4 (S100A4) expression (Figure 4C).

**Figure 4.**
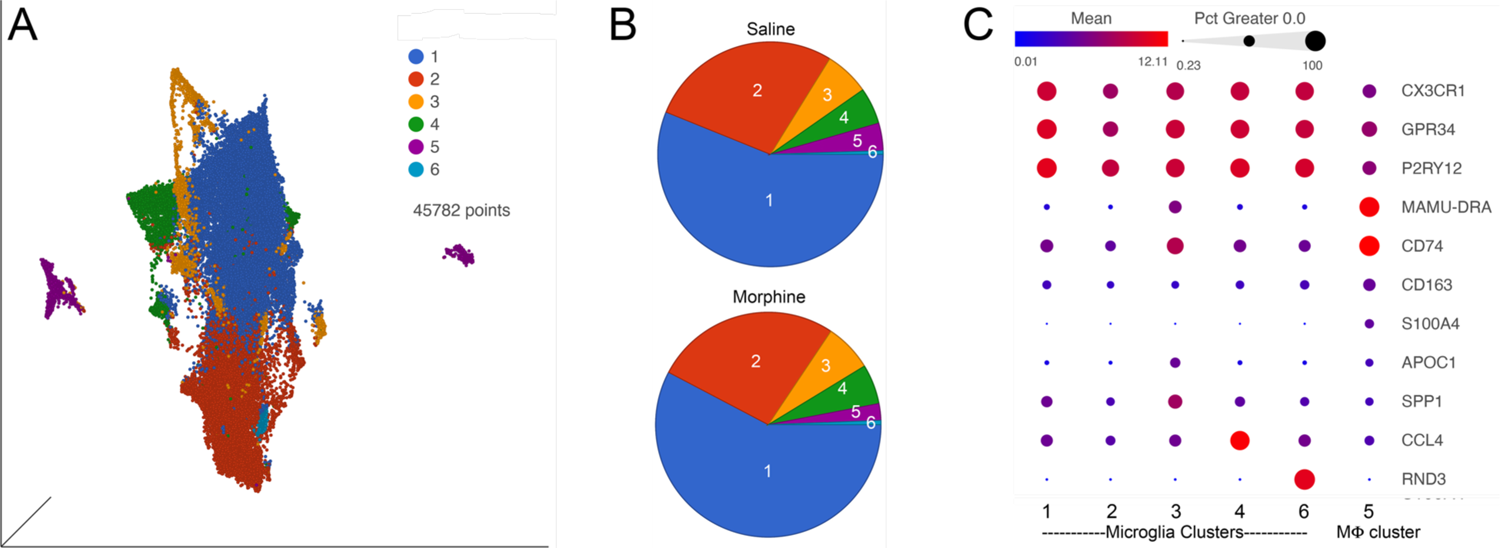
ScRNA-seq analysis of brain macrophages and microglia from the saline and morphine groups of animals. **A)** Uniform Manifold Approximation and Projection (UMAP) representation of the cells, grouped in 6 clusters and differentially colored as determined by graph-based clustering. **B)** Pie charts of the proportion of cells in each cluster for the saline and morphine groups. **C)** Bubble plot of selected genes differentiating the clusters. The color indicates the least-square mean expression (log_2_), and the size of the circle indicates the percentage of cells in which gene expression was detected.

Expression values for the cells in the clusters were examined for differentially expressed genes (DEGs) versus the other clusters, some of the genes are illustrated in Figure 4C. Clusters 1 and 2 did not differ much in DEGs compared to the other clusters, and showed some differences with each other, with complement C1q B chain (C1QB) and beta-2-microglobulin (B2M) higher in cluster 1, and activating transcription factor 3 (ATF3) and FosB proto-oncogene, AP-1 transcription factor subunit (FOSB) higher in cluster 2. Cluster 3 was notable for higher levels of numerous genes, including increases in apolipoprotein C1 (APOC1), major histocompatibility complex, class II, DR alpha (MAMU-DRA), and secreted phosphoprotein 1 (SPP1). Cluster 4 had increased expression of C-C motif chemokine 3-like and 4 (CCL3L and CCL4), whereas in cluster 6 Rho family GTPase 3 (RND3) was greatly increased. In general clusters 1 and 2 expressed a homeostatic microglia pattern (71–75). The differences between cluster 1 and 2 may be stress-related, as ATF3 and FOSB are members of the AP-1 transcription factor family, induced in acute stress (76). ATF3 has been proposed to be induced as a negative-feedback loop to regulate macrophage inflammatory cytokine production (77). APOC1 and SPP1, prominent in cluster 3, are two of the genes characterizing the DAM subtype of microglia found in animal models of Alzheimer’s disease and other conditions (72, 78). Regarding cluster 4, CCL4 (as well as CCL3) expression characterizes a small population of microglia that increase with aging in mice (71). The high expression of RND3 is intriguing as it has been proposed to induce a neuroprotective phenotype in microglia (79). A complete list of DEGs between the clusters is presented in Supplemental Table 2.

### Chronic morphine alters CNS myeloid cell gene expression leading to a neurodegeneration-related phenotype

We next examined the clusters for DEGs in the context of morphine administration. The microglial clusters showed appreciable similarity in the DEGs. In the five microglia clusters (clusters 1-4 and 6), 17 genes were upregulated in common. Microglia cluster 6 failed to exhibit morphine downregulated genes. Excluding cluster 6, there were 33 upregulated, and 22 downregulated, genes that were common in microglia clusters 1-4 (Supplemental Figure 2).

We used gene set enrichment analysis (GSEA) to assess whether morphine altered the phenotype of the clusters in a characteristic manner. We used curated gene lists (Supplemental Table 4) for disease-associated (DAM) (72) and activation-response (ARM) (73) microglia, as well as neurodegeneration-related brain myeloid cells (78). As shown in Table 1, morphine induced a neurodegeneration-related myeloid cell phenotype in all the microglia clusters except the smallest cluster, cluster 6, and a DAM and ARM phenotype in the largest cluster, cluster 1. Macrophage cluster 5 also took on a neurodegeneration-related phenotype in response to morphine.

**Table 1.** Gene set enrichment analysis. The enrichment score (ES) is the degree to which a set is overrepresented comparing the morphine with the saline group. The normalized enrichment score (NES) indicates the degree to which a gene set is overrepresented for differences in gene set size, normalized for correlations between gene sets and the expression dataset. The positive numbers connote enrichment in the morphine DEGs for that cluster.

| Cluster # | Gene Set | p-value | FDR q-value | ES | NES |
| --- | --- | --- | --- | --- | --- |
| **Microglia Clusters** |  |  |  |  |  |
| 1 | Neurodegeneration-related | 0.003 | 0.006 | 0.39 | 2.18 |
| 2 | Neurodegeneration-related | 0.008 | 0.03 | 0.32 | 1.9 |
| 3 | Neurodegeneration-related | 0 | 0.001 | 0.44 | 2.21 |
| 4 | Neurodegeneration-related | 0.012 | 0.037 | 0.41 | 1.82 |
| 1 | Activation response | 0.012 | 0.025 | 0.34 | 1.87 |
| 1 | Disease-associated | 0.037 | 0.04 | 0.22 | 1.64 |
| **Macrophage Cluster** |  |  |  |  |  |
| 5 | Neurodegeneration-related | 0.006 | 0.025 | 0.47 | 1.82 |

The DEGs from the morphine versus saline condition in each cluster were then used for Ingenuity Pathway Analysis (IPA). We focused on the canonical pathways, and the results are presented in Supplemental Table 5; selected pathways are shown in Table 2. Interestingly the phagosome formation pathway was increased in 3 of the microglia clusters, and several signaling pathways were increased in the microglia clusters, including those of neuroinflammation in cluster 3, TREM1 in cluster 4, and G-protein coupled receptor in cluster 6. Signaling pathways were also enriched in macrophage cluster 5, including those involved in leukocyte extravasation and neuroinflammation.

**Table 2.** IPA canonical pathways. Selected predicted pathways with significance (p-value < 0.05) and increased activation (Z-score > 1), as determined using the DEGs between cells from morphine and saline groups within each cluster. PPR = Pattern Recognition Receptors.

| Cluster # | Ingenuity Canonical Pathways | p-value | Ratio | z-score |
| --- | --- | --- | --- | --- |
| **Microglia Clusters** |  |  |  |  |
| 2 | Phagosome Formation | 0.009 | 0.015 | 1.897 |
| 3 | Neuroinflammation Signaling Pathway | 0.038 | 0.019 | 2 |
| 3 | cAMP-mediated signaling | 0.037 | 0.021 | 1.342 |
| 4 | Phagosome Formation | 0.004 | 0.022 | 2.324 |
| 4 | Calcium Signaling | 0.021 | 0.028 | 2.236 |
| 4 | Senescence Pathway | 0.010 | 0.027 | 2.121 |
| 4 | Role of PPR in Recognition of Bacteria & Viruses | 0.005 | 0.039 | 2 |
| 4 | TREM1 Signaling | 0.007 | 0.052 | 2 |
| 6 | Phagosome Formation | 0.015 | 0.013 | 3 |
| 6 | G-Protein Coupled Receptor Signaling | 0.042 | 0.011 | 2.828 |
| 6 | ERK/MAPK Signaling | 0.032 | 0.019 | 2 |
| **Macrophage Cluster** |  |  |  |  |
| 5 | PPARα/RXRα Activation | 0.048 | 0.036 | 2.236 |
| 5 | Leukocyte Extravasation Signaling | 0.017 | 0.042 | 1.633 |
| 5 | RHOA Signaling | 0.019 | 0.048 | 1.633 |
| 5 | PTEN Signaling | 0.014 | 0.047 | 1.342 |
| 5 | Integrin Signaling | 0.029 | 0.038 | 1.342 |
| 5 | PPAR Signaling | 0.035 | 0.047 | 1.342 |
| 5 | Neuroinflammation Signaling Pathway | 0 | 0.057 | 1.213 |
| 5 | Necroptosis Signaling Pathway | 0.018 | 0.045 | 1.134 |

Several morphine-upregulated DEGs are illustrated in Figure 5. In the microglia clusters 1-4 and 6, transcripts for immediate early response 3 (IER3), a feedback inhibitor of NF-κB in the inflammatory response (80), were increased, as was the mRNA for receptor activity-modifying protein 1 (RAMP1), a component of the calcitonin gene-related peptide receptor complex, which has microglia-activating activity (81). Also increased was the sodium bicarbonate cotransporter 3 (SLC4A7), critical for phagosome acidification (82), and SPP1 (especially prominent in cluster 3), a pleiotropic cytokine with numerous links to CNS diseases (83). There was no such change in the expression of these genes in macrophage cluster 5. Instead, several genes were upregulated in the morphine group, including C-type lectin domain family 7 member A (CLEC7A), a myeloid-predominant pattern-recognition receptor (84), type II major histocompatibility genes such as MAMU-DQA1, which are increased in macrophage activation (85), and the intermediate filament protein gene vimentin (VIM), which is increased in macrophages differentiation and activation (86). None of these genes had appreciable expression in the microglial clusters.

**Figure 5.**
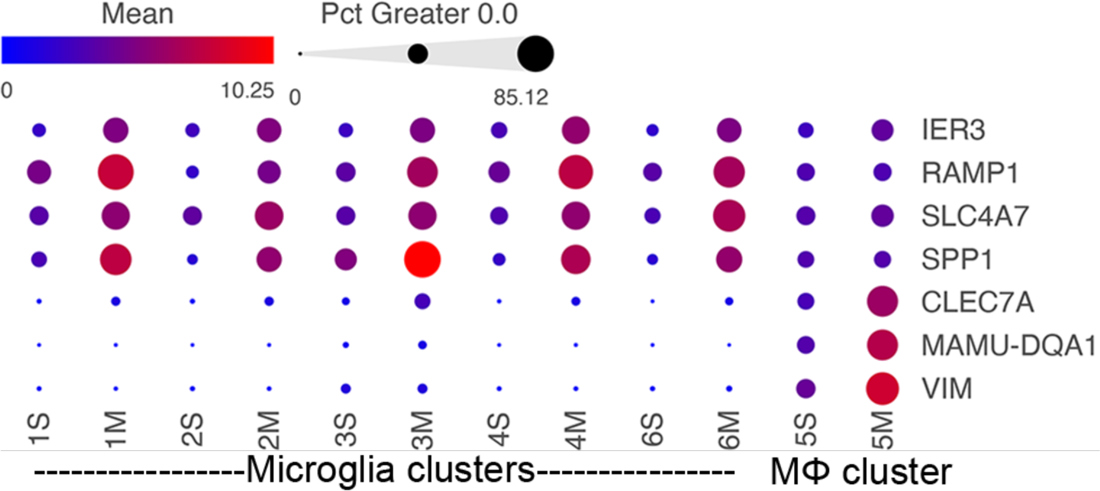
Morphine-related DEGs expression in the different clusters. Bubble plot of selected DEGs between the saline and morphine cells within the clusters. The color indicates the least-square mean expression (log_2_), and the size of the circle indicates the percentage of cells in which gene expression was detected.

### Osteopontin expression is increased in microglia, particularly in white matter microglia, by morphine in SIV-infected monkeys

In examining the DEGs, we were particularly intrigued by the increase in SPP1 (also known as osteopontin), a pleiotropic molecule produced by, and with numerous effects on, macrophage and microglia, as well as with links to neurodegenerative disorders and many other diseases (83, 87–91). Its expression was significantly increased in all microglia clusters, but not in the macrophage cluster (Table 3).

**Table 3.** Morphine effect on SPP1 expression in the brain macrophage and microglia clusters. The false discovery rate (FDR) q-value is given for the comparison of the expression levels in cells from the saline and morphine groups. The fold-change in expression is listed, based on the least-square (LS) means of the cells in each group.

| **Microglia cluster #** | **FDR**<br>**q-value** | **SPP1**<br>**Fold-change** | **Saline**<br>**LS-Mean** | **Morphine**<br>**LS-Mean** |
| --- | --- | --- | --- | --- |
| **1** | 0 | 4.28 | 417.49 | 1785.87 |
| **2** | 0 | 4.51 | 257.37 | 1160.59 |
| **3** | 2.71E-182 | 3.22 | 2618.24 | 8419.98 |
| **4** | 3.51E-134 | 5.29 | 277.74 | 1470.48 |
| **6** | 3.03E-08 | 3.43 | 316.23 | 1085.12 |
| **Macrophage cluster 5** | 0.795 | 1.11 | 561.65 | 623.99 |

SPP1/osteopontin is known to be expressed by neurons, oligodendrocytes, and microglia, as well as CNS-infiltrating immune cells, with different patterns in different brain regions, and can change in reaction to stimuli (92–97). To address regional and cellular expression, we performed single-nucleus RNA sequencing (snRNA-seq) on two brain regions – frontal lobe white matter and the caudate nucleus. Again, four animals from each group were examined. An average of ∼5,600 nuclei were examined from each animal from the white matter and ∼5,900 nuclei from the caudate. Clustering and marker gene analysis of the white matter nuclei revealed 1 cluster of oligodendrocyte precursor cells (OPC), 3 clusters of oligodendrocytes, 1 cluster of mixed astrocytes and neurons, and 1 cluster of microglia (containing 4,761 nuclei, 10% of all nuclei) (Figure 6A). In the caudate, 1 cluster of OPC, 3 clusters of oligodendrocytes, 1 cluster of astrocytes, 3 clusters of neurons, and 1 cluster of microglia (containing 3,806 nuclei, 8.1% of all nuclei) were identified (Figure 76). In both, cluster 4 contained microglia (P2RY12 and CSF1R positive).

**Figure 6.**
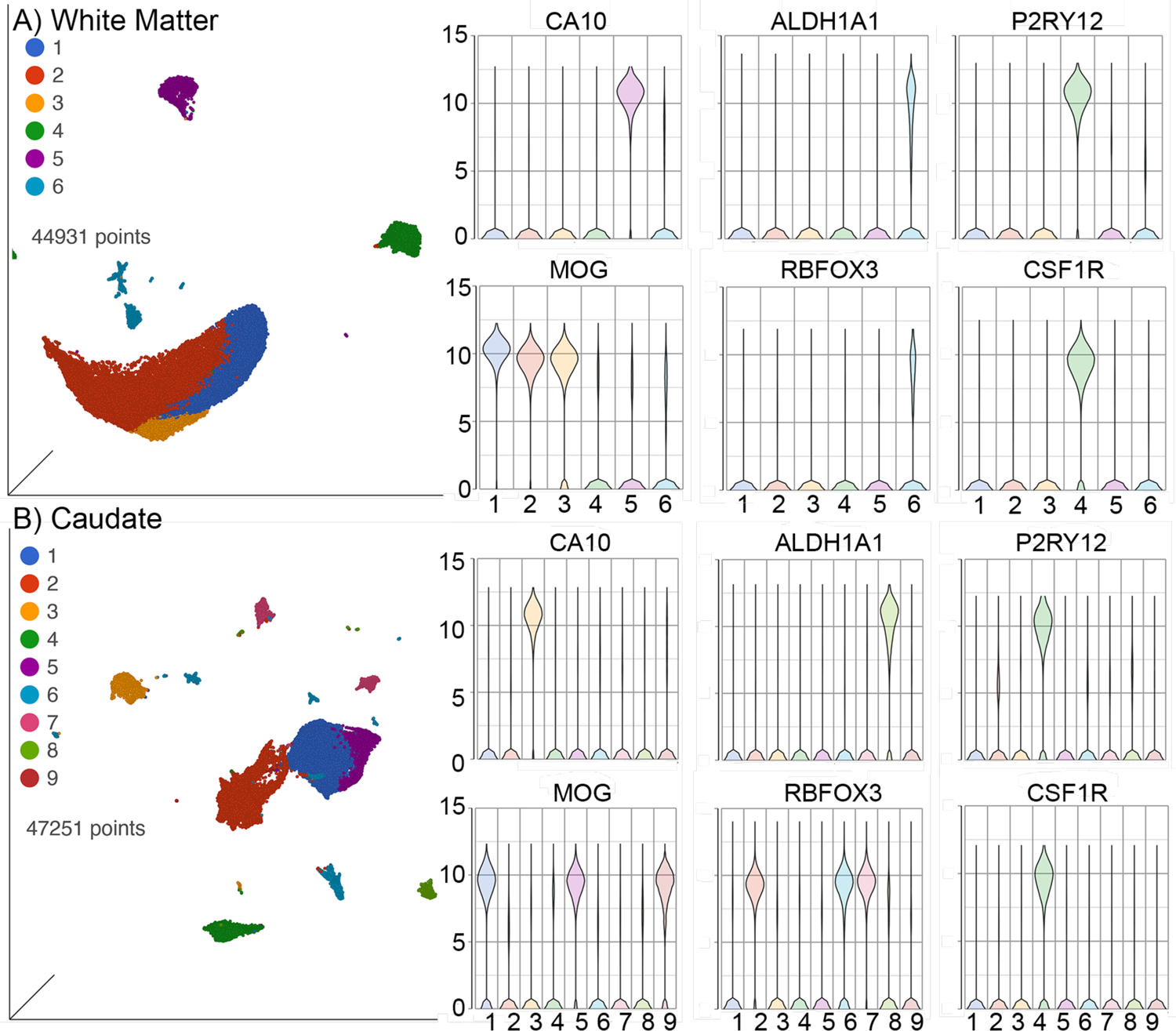
SnRNA-seq of brain regions. A) White Matter and B) Caudate. Left in each are UMAP plots, right are violin plots of cellular identifying markers. OPC express carbonic anhydrase 10 (CA10), oligodendrocytes myelin oligodendrocyte glycoprotein (MOG), astrocytes aldehyde dehydrogenase 1 family member A1 (ALDH1A1), neurons RNA binding Fox-1 homolog 3 (RBFOX3, also known as neuronal nuclei, NeuN), and microglia purinergic receptor P2Y12 (P2RY12) and colony-stimulating factor-1 receptor (CSF1R). Values on the Y-axis are the least-square mean expression (log_2_).

Examination of SPP1 expression revealed that in the white matter (Figure 7A), the highest expression was found in the microglia (cluster 4), with most of the oligodendrocytes in clusters 1 and 2 expressing SPP1, fewer in oligodendrocyte cluster 3, and still fewer in OPC cluster 5 and neuron/astrocyte cluster 6. In the caudate (Figure 7B), expression was predominantly in two of the oligodendrocyte clusters, clusters 1 and 5, with low-level expression in cells in microglia cluster 4 and the other cellular clusters. The white matter microglia had a much higher level of expression of SPP1 than caudate microglia, and morphine increased SPP1 expression only in the white matter microglia (Table 4, Figure 7C). Although at a much lower expression level, in the caudate, morphine conversely lowered SPP1 expression in microglia, as it did in the oligodendrocyte clusters (Table 4).

**Figure 7.**
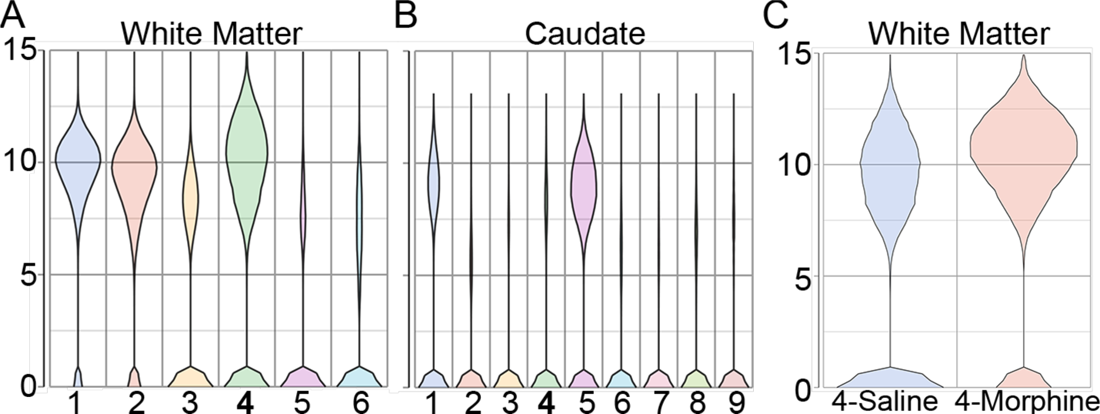
Violin plots of SPP1 expression from snRNA-seq. **A,B)** White Matter and Caudate. The microglia cluster (cluster 4 in both) is indicated in bold. **C)** Expression in cluster 4, containing microglia from the white matter, in the saline and morphine groups. Values on the Y-axis are the least-square mean expression (log_2_).

**Table 4.** Morphine effect on SPP1 expression in nuclei from the white matter and caudate. The false discovery rate (FDR) q-value is given for the comparison of the expression levels in cells from the saline group and the morphine group. The fold change in expression is listed, based on the least-square (LS) means of the cells in each group. Only microglia clusters and those with appreciable SPP1 expression are shown.

| **Site/Cluster** | **Cell type** | **FDR**<br>**q-value** | **SPP1**<br>**Fold-change** | **Saline**<br>**LS-Mean** | **Morphine**<br>**LS-Mean** |
| --- | --- | --- | --- | --- | --- |
| White Matter/4 | Microglia | 4.89E-131 | 1.95 | 1126.06 | 2200.74 |
| Caudate/4 | Microglia | 1.28E-04 | -1.68 | 131.47 | 78.40 |
| White Matter/1 | Oligodendrocyte | 6.58E-18 | -1.24 | 1198.42 | 970.20 |
| White Matter/2 | Oligodendrocyte | 2.12E-18 | -1.33 | 949.81 | 713.96 |
| White Matter/3 | Oligodendrocyte | 2.28E-34 | -1.39 | 278.53 | 200.97 |
| Caudate/1 | Oligodendrocyte | 1.04E-299 | -2.46 | 628.53 | 255.62 |
| Caudate/5 | Oligodendrocyte | 1.22E-51 | -2.77 | 675.22 | 243.93 |

## DISCUSSION

Substance use disorders, and more specifically abuse of opioids, are important comorbidities among PWH. The current study investigated interactive effects of opioids in the brains of SIV-infected rhesus macaques with suppressed viremia. The primary objective was to examine factors that can contribute to neuroHIV, including the increased viral reservoir found in morphine-treated animals. While HIV encephalitis dominated neuropathogenesis in the pre-treatment era, in the current era of efficacious treatment neuropathological studies have not uncovered the substrate of central nervous system (CNS) dysfunction (98, 99). While ample studies have focused on the characterization of latent virus reservoirs in the peripheral circulating CD4+ T cells and lymphoid tissues, there is a paucity of information on the mechanisms involved in the persistence of HIV reservoirs in the myeloid cells of the CNS.

To assess the impact of infection and cART in the context of morphine dependence, we evaluated the expression of biomarkers as well as lymphocyte subsets before infection, during the acute phase of infection at peak viremia (2 weeks p.i.), post-peak viremia (5 weeks p.i.) as well as at two additional time points post-virus suppression following cART (22 and 35 weeks p.i., which correspond to 17- and 30-weeks post-cART initiation, respectively). Before infection, morphine exposure itself led to an anti-inflammatory phenotype. For example, the expression of adiponectin, a hormone known to attenuate the inflammatory response, was significantly upregulated in the presence of morphine. Further, expression of several other inflammatory mediators and markers such as IL-16, MMP-2, TNFR2, and VCAM-1 were all downregulated in the plasma of uninfected morphine-dependent macaques relative to uninfected saline-treated animals.

Striking results were found during the acute infection phase, which is crucial in determining disease course in untreated infection. The increase in CD8+ T cells and NK cells, characteristic of response to infection in the saline group, was largely attenuated in the morphine group. Expression of CRP, a liver-produced marker of inflammation, increased dramatically in saline-treated SIV-infected animals, while its levels remained unchanged over baseline in the presence of morphine. Similarly, ferritin, an iron-sequestering protein that is elevated during inflammation, increased significantly in the saline versus morphine-dependent macaques. IL-1RA, IL-16, IL-18, MMP2, TNFR2, and VCAM-1 were all significantly lower at this acute time point in the morphine group compared with the saline group. These findings are in keeping with published reports implicating attenuation of the inflammatory immune response by morphine in the periphery of SIV-infected macaques (41). Along these lines, heroin abusers have been shown to exhibit reduced immune responses and increased incidence of infections (100, 101). Furthermore, chronic administration of relatively low doses of morphine in a cohort of rhesus monkeys over a 2-year period was shown to affect immunocompetence, suppressing NK-cell activity and decreasing the percentage of CD4+ circulating lymphocytes (102, 103). In addition a study on morphine-treated SIV-infected monkeys found that they also showed depressed weight gain as well as were more vulnerable to opportunistic infections with quicker progression to simian AIDS (104). Studies in a variety of systems confirm the immune suppressive properties of opioids (105).

Most of the effects of morphine found in the uninfected animals persisted even following cART in the infected, morphine-dependent animals compared with the infected saline group. It is likely that a muted inflammatory response during the initial infection period could likely contribute to increased seeding of virus reservoirs with decreased clearance of cells resistant to the cytotoxic effects of the virus, such as macrophages and microglia. We did not see any change in antiretroviral concentrations in either brain, or biofluids in the context of morphine, likely indicating that drug pharmacokinetics played a negligible role in the maintenance of CNS virus reservoirs in our model system.

Brain-resident myeloid cells – macrophages and microglia – are the primary HIV/SIV infected cells in the brain and comprise the brain reservoir during treatment (48, 49). These cells play multifaceted roles and can control CNS damage owing to their ability to elicit key immune defenses in the CNS and their inherent ability to release proinflammatory cytokines and chemokines in response to cellular insults (106, 107). In the CNS, the deleterious effects of infection include the generation of proinflammatory mediators within these infected cells, which then in turn cause bystander dysfunction and toxicity in uninfected neurons and glia. Drugs of abuse, including opioids such as morphine, also have distinct effects on brain microglia and macrophages.

To assess the effects of morphine on microglia and macrophages, we performed scRNA-seq on brain cells from the two groups of monkeys. The proportion of different microglia subtypes, identified by clustering and proportion of macrophages, did not differ between the groups. Both the microglia and the macrophages showed changes in gene expression between the saline-administered and morphine-dependent animals. While some differences were present between the microglia subtypes, the upregulated and downregulated genes showed commonalities. In the morphine-treated group, expression of neurodegeneration-related genes was increased in all microglia subtypes and macrophages. Enrichment of genes related to the DAM phenotype was present in one of the microglia clusters due to morphine. However, no specific DAM cluster of microglia was identified in the morphine group. While the DAM phenotype is prominent in mouse models of neurodegeneration, its translation to humans is inconsistent, and thus calls into question whether a specific DAM microglia cluster could be found in NHP. Pathway analysis was notable for increases in the phagosome formation pathway in microglia as well as the neuroinflammation signaling pathway in microglia and macrophages, as well as other signaling pathways in the different cellular clusters.

The DEGs in context of morphine were different in the macrophage cluster than those in the microglia clusters. In microglia clusters, morphine increased the expression of SPP1/osteopontin, which has been linked to HIV-associated neurocognitive disorders in several studies. While our microglia preparations for the scRNA-seq were derived from the whole brain, to obtain an independent assessment of morphine-related increase in SPP1 transcripts in microglia, we performed snRNA-seq on nuclei isolated from two brain regions, frontal lobe white matter and the caudate. In both regions we could identify a cluster of cells that represented microglia. Somewhat surprisingly there was an anatomic location difference in SPP1 expression, both for the value for SPP1 expression in the saline group as well as the magnitude and direction of change in the morphine group. Much higher levels of SPP1 were found in white matter microglia than in caudate microglia in the saline animals. In the morphine group the SPP1 expression level was significantly increased in the white matter, whereas in caudate microglia SPP1 expression was lower. Interestingly a study on human brains found higher SPP1 expression in white matter versus grey matter microglia, and the difference increased with aging (108). Another study in mice found distinct clusters of microglia only within the white matter, denoted as white matter-associated microglia (WAMs), and such clusters were enriched SPP1 expression (109). This study found that WAMs increased with aging and expressed a subset of the DAM gene signature identified in Alzheimer’s disease models. In addition to WAMs, other differences reported between white and grey matter microglia include a higher density of microglia in the white matter and increased markers of antigen presentation and phagocytosis (110). Numerous abnormalities in white matter have been characterized in PWH and some are found to be associated with neurocognitive disorders (111, 112). Interestingly, opioids have also been shown to induce white matter damage (113).

The role of SPP1/osteopontin in HIV infection has been assessed in several studies. We first reported increased osteopontin mRNA expression in the brains of SIV-infected rhesus monkeys with encephalitis (114), and that osteopontin protein was increased in the plasma and CSF of animals with SIV encephalitis (115). We subsequently found increased osteopontin levels in the CSF of PWH and in the plasma of PWH with dementia (116). Others have also found correlations with osteopontin. In PWH, regardless of cART treatment, osteopontin was found to be elevated in the plasma (117). In the basal ganglia of SIV-infected monkeys, osteopontin mRNA was highly expressed compared to that in the uninfected animals or those with viral suppression by cART (118). In another study in PWH, osteopontin was found to be increased in the brain and CSF, and importantly, shown to enhance HIV replication in primary human macrophages (119, 120). Further, osteopontin can affect monocyte entry into, and egress from, the brain. Osteopontin is a chemotactic agent for macrophages and increases their survival (121–123). In addition, we found that osteopontin prevents monocyte recirculation after traversing a barrier, and reduces monocyte apoptosis, thus retaining monocytes that enter the brain (115). SPP1 is frequently found in gene lists of activated or disease-linked microglia subsets identified by scRNA-seq in mice and humans (124). Interestingly in mice, a microglia subset expressing SPP1/osteopontin has been found to execute proinflammatory responses (125).

Osteopontin actions are also regulated through several mechanisms. Alternate translational start sites lead to an intracellular and secreted forms (126). The intracellular form predominates in myeloid cells, and has distinct properties compared to the secreted form, which is abundantly produced in T cells (127). Secreted osteopontin can be cleaved by thrombin, leading to enhanced chemotactic activity on monocytes (128). Furthermore, genetic polymorphisms exist in the osteopontin gene, and alternate splicing can produces different isoforms of osteopontin (127). Post-translational phosphorylation also leads to differential activities on macrophage migration and macrophage activation (129). The role of these different forms of osteopontin, and their effect on brain microglia, macrophages, HIV, and the effects of drugs of abuse such as opioids remains a significant gap in our knowledge and awaits future study.

Using scRNA-seq of microglia from SIV-infected animals, we recently found that in untreated animals, microglia exhibited a proinflammatory state, and that viral suppression with cART restored the microglia to a homeostatic-like state (130). Here we find that in the setting of viral suppression with cART, morphine had numerous other effects on the microglia as well as brain macrophages. Strikingly, the 4 largest microglia clusters and the macrophage cluster all showed enriched expression of genes related to neurodegeneration. These genes were curated as a core set of genes exhibiting increased expression in CNS myeloid cells in multiple models of neurodegenerative disease. Interestingly, it was suggested that in neurodegeneration this increase in expression changes how these cells interact with their environment, since the majority of the genes in this classification were annotated with a Gene Ontology (GO) term associated with either the plasma membrane or extracellular space (78). While we did not find separate clusters of CNS myeloid cells in the morphine-treated group samples, all clusters of these cells were enriched for increased expression of neurodegeneration-related genes.

The enrichment for the DAM phenotype in one of the clusters of microglia and macrophages reveals a similar response to damage found in a variety of experimental animal models of CNS disease, and, to an extent, in human disease. In addition, we found that administration of morphine increased the expression levels of genes regulating the neuroinflammation signaling pathway in macrophages. Other myeloid signaling pathways, such as the integrin and triggering receptor expressed on myeloid cells 1 (TREM1) pathway, were also increased different myeloid clusters and the macrophage cluster. These, along with an increase in the phagosome formation pathway in 3 clusters, connote responsiveness of the microglia and macrophages to morphine within the brains of SIV-infected NHP. While there is evidence of role of TREM1 in resistance to HIV-mediated cytopathogenesis in macrophages (131, 132), this effect, however, has not been examined thus far in the context of opioids.

Very few studies using scRNA-seq for assessing the opioid effects have been performed. In one study, cells from the nucleus accumbens (NAc) were studied following acute morphine treatment in mice. Four hours after morphine administration, a cluster of dopamine receptor 1a expressing neurons, astrocytes, and multiple clusters of oligodendrocytes, including OPC, were found to respond transcriptionally to the drug, with the most robust response found in the oligodendrocytes (133). While we did not examine the NAc, we note that we did not see similar responses in the oligodendrocytes from the two regions we examined in the brain, including the caudate nucleus, also part of the striatum. Additionally, our model was of chronic, not acute, morphine treatment. Another study examined peripheral blood mononuclear cells from opioid-dependent and control subjects (134). Here, in circulating naïve monocytes, suppression of antiviral genes was found, and the response to stimulation in monocytes and other blood cell types was suppressed. *In vitro* morphine treatment of blood immune cells yielded similar results. We indeed found evidence for such morphine-linked immunosuppression throughout the longitudinal course of infection in the morphine-treated group.

ScRNA-seq CNS studies in the context of HIV infection have also been limited, in part, understandably, to the inaccessibility of the brain during life. However the CSF has been studied and interesting results have been found. The first study examined the CSF cells from PWH virally suppressed on cART compared to uninfected controls and identified a small population of myeloid cells in the PWH subjects with increased expression of genes found in microglia and specifically in neurodegeneration-associated microglia clusters (135). This group then expanded these studies, and again identifying this rare cluster of microglia-like cells in the CSF, as well as finding CD4+ T cells expressing HIV in the CSF from these virally-suppressed subjects (136). Parallel examination of the CSF cells and the brain in SIV-infected rhesus macaques could help clarify the role of this unique subset of myeloid cells in the CSF in HIV/SIV infection, as well as any effect of opioids.

Limitations to our study include the lack of analysis of uninfected animals, in both control and morphine groups, and a relatively small sample size. Only male animals were used in this study, making it difficult to make any inferences on sex-specific differences. Furthermore, in our study, the length of morphine-dependence and duration of viral infection (which was treated early to suppress viremia) was much shorter than that observed in PWH. Numerous other molecules which could potentially affect the HIV reservoirs were differentially regulated by morphine in the brain macrophages and microglia and were not investigated further other than through GSEA and pathway analysis. We could not discern the effects of different forms of osteopontin analysis on the brain reservoir in this model. Other than brain macrophages and microglia, the actions of morphine on other cell types in the brain were not examined.

In summary, morphine, a prototypic opioid, exhibited marked effects on NHP before and after SIV infection. A general immune suppressive environment was found based on immune cell and soluble mediator analysis in the blood. Following sustained viral suppression, we analyzed brain macrophages and microglia, targets for SIV and HIV, and mediators of both protective and destructive actions as well as the CNS reservoir for these viruses by scRNA-seq and snRNA-seq. These findings revealed gene expression changes that could indicate transition to a neurodegenerative phenotype due to morphine administration, and changes that likely contribute to the basis of our prior findings of increased viral reservoirs in the brains of cART-suppressed, morphine-dependent animals. In summary, the overall immune suppressive environment, altered gene expression by CNS myeloid cells, and the actions of one of the morphine-induced genes, SPP1/osteopontin, yield novel insights into the interactions of the dual-linked health crises of HIV and opioids.

## METHODS

### Animals

The animals used in this study have been previously described (57). Briefly, the rhesus macaques tested negative for a panel of viral and bacterial pathogens before their use in the study. Animals in the morphine group received intramuscular injections of morphine, which were ramped-up over a 2-week period to a final dose of 6 mg/kg morphine administered twice daily on weekdays, and once daily on weekends. This dose was maintained for 7 weeks. Animals in the saline group received saline injections on the same schedule. Animals were then inoculated with SIVmac251. Five-weeks post-inoculation, all animals received a once-daily subcutaneous injection of 1 mL/kg of body weight containing 40 mg/mL emtricitabine (FTC), 20 mg/mL tenofovir (TDF) and 2.5 mg/mL dolutegravir (DTG), continued until the time of necropsy. At necropsy, blood-borne cells were cleared by perfusion with PBS containing 1U/mL heparin. Microglia/macrophage-enriched cellular isolation from the brain was performed using our previously described procedure of physical and enzymatic disassociation followed by Ficoll gradient purification (10, 137), and then cryopreserved in the presence of 10% DMSO / 90% fetal bovine serum and stored over liquid nitrogen. Brain regions were dissected by a board-certified pathologist, frozen, and stored at −80° C.

### Blood cell subsets, Albumin, and Flow cytometry

Complete blood counts (CBC) and chemistry/metabolic panels were performed in the CLIA-certified Pathology labs of Nebraska Medicine (Omaha, NE). Overall cell numbers and proportions of the general classes of cells were reported on the CBC; the albumin concentration was determined on the chemistry/metabolic panel.

Flow cytometry evaluation was completed on peripheral whole blood and assessed prior to viral inoculation and longitudinally throughout the study. Cells and staining reagents were kept on ice, centrifugation was conducted at 4° C and 3500 × g. Cells were first stained with UV blue Live dead assay (Invitrogen, Carlsbad, CA), washed, blocked using flow cytometry buffer (eBioscience, Waltham, MA), pelleted, and surface antigens stained with a cocktail containing brilliant stain buffer (BD Biosciences, San Jose, CA) and fluorochrome-conjugated monoclonal antibodies against CD11b-BV605 (Biolegend, San Diego, CA), CD14-ECD (Beckman Coulter, Brea CA), CD3-BV421, CD4-BV786, CD8-PeCy7, CD16-APCH7, and CD20-BV711 (all from BD Bioscience). Stained whole blood was washed with flow cytometry buffer, pelleted and red blood cells lysed with red blood cell lysis buffer (Sigma Aldrich, St. Louis, MO). Lysed samples were washed in flow cytometry buffer, pelleted, and fixed using 1% paraformaldehyde in flow cytometry buffer. Cells were stored overnight at 4° C before acquisition. All events were acquired using the BD Fortessa X450, and data were analyzed using BD FACSDiva v9.0 or FlowJo for Mac version 10.6 (BD Biosciences).

The lymphocyte percentage was determined as proportion of white blood cells measured on the day of draw CBC. Lymphocyte percentage was multiplied by the white blood cell count to obtain absolute lymphocyte count per microliter, and further multiplied by the flow cytometric determination of the CD3+CD4+ and CD3+CD8+ proportions within the lymphocyte FSC/SSC gate on live cells to obtain absolute number of CD4+ and CD8+ T-cells, respectively, per microliter of blood.

### Plasma biomarkers

Plasma, collected in EDTA-anticoagulated blood, was separated within an hour of collection and stored frozen at −80° C. Samples were sent to Myriad RBM (Austin, TX), a CLIA-certified lab, for multiplexed biomarker assay using the Human v2.0 Multianalyte profile (MAP) immunoassay. We previously reported the normal range for the 46 analytes that can be detected in rhesus macaques (138).

### Antiretroviral quantitation

Concentrations of TFV, FTC, and DTG in plasma and CSF, and DTG and the intracellular triphosphates (TP), tenofovir-diphosphate (TFV-DP) and emtricitabine-triphosphate (FTC-TP) in tissue homogenates, were quantified by liquid chromatography with tandem mass spectrometry with previously described validated methods (139–141).

### ScRNA-seq / snRNA-seq

From the 11 SIV-infected, cART-treated animals (6 in the morphine group, 5 in the saline group), 4 from each group were used for the single cell/nucleus analyses. For scRNA-seq of microglia, the saline group consisted of samples from animals 74T (referred to as CI47 in (57)), 77T (CK40), 82T (CI04), and 83T (CK03), and the morphine group samples from animals 73T (CK22), 75T (CK48), 78T (CI90), and 80T (CH89). The scRNA-seq data for the three of the members of the saline group, 74T, 82T, and 83T, were reported previously (130). For the snRNA-seq several animal samples were substituted within the groups due to the availability or suitability of samples for processing and analysis. For the frontal lobe white matter, 81T (CK07) was substituted for 82T for the saline group and 76T (CK31) for 80T in the morphine group. For the caudate those same substitutions were made, in addition morphine 79T (CI83) was substituted for 75T in the morphine group.

scRNA-seq procedures were completed using our previously described methods (137, 142). Briefly, cryopreserved microglia were treated with 1% DNase (Sigma Aldrich), washed, stained with UV-blue live/dead assay (Invitrogen, Waltham, MA). Post stain, microglia were washed, resuspended in MACS separation buffer with 0.1% bovine serum albumin, isolates were then enriched with nonhuman primate anti-CD11b microbeads (Miltenyi, Gladbach, Germany), followed by washing and positive selection on MACS Separator LS columns. Recovered cells were stained anti-mouse/human CD11b antibody clone M1/70 (BV605; Biolegend, San Diego, CA), washed, and sorted based on size, singlets, live, CD11b+ events using an Aria2 flow cytometer (BD Biosciences, San Diego, CA).

For snRNA-seq, caudate tissue was chopped fine, transferred to a Dounce homogenizer with 300 µL of NP40 lysis buffer (consisting of 10 mM Tris-HCL (pH 7.4), 10 mM NaCl, 3 mM MgCl_2_, 1% nonidet P40 substitute, 1 mM DTT, 1 U/µL RNase inhibitor; all from Sigma Aldrich, St. Louis, MO). Samples were homogenized, rested on wet ice, washed in 1% BSA in dPBS, rested on wet ice, and filtered through 70 µm filter. Samples were volume adjusted to 1 mL and stained with 7-AAD staining solution (Sigma Aldrich). Nuclear events were captured based on size, singlets, and nuclear based on 7-AAD stain, using an Aria2 flow cytometer.

A similar method was used to isolate nuclei from white matter samples, except after filtration samples were adjusted to a total volume of 2 mL of 1% BSA in dPBS. Samples were divided into two 2 mL tubes. In each tube, 1 mL of filtered sample was mixed with 300 µL of debris removal solution (Miltenyi) overlaid with 1 mL 1% BSA in dPBS. Samples were centrifuged at 3000 x G for 10 minutes. Subsequent upper layers were discarded and lower layer, containing the nuclei, were washed in ice cold 1% BSA in dPBS. Nuclei isolates were washed again, filtered through 70 µm filter and final volume was adjusted to 1 mL. Samples were then stained with 7-AAD and sorted on an Aria 2 flow cytometer.

Post-processing, isolates were concentrated to approximately 1,000 cells or nuclei/μL and assessed by trypan blue for viability and concentration. Based on 10x Genomics parameters targeting 8000 cells, the calculated volume of cells or nuclei was loaded onto the 10x Genomics (Pleasanton, CA) Chromium GEM Chip and placed into Chromium Controller for cell capturing and library preparation. We used the 10x Genomics Single Cell 3’ GEM, Library, and Gel bead kit v3.1. The prepared libraries were sequenced using a NovaSeq6000 sequencer (Illumina, San Diego, CA) using Kit S1-100 Version 1.5 for scRNA-seq and Kit S2-100 Version 1.5 for snRNA-seq.

### Bioinformatics

We generated FASTQ files and count matrices using 10x Genomics cellranger pipeline version 3.1.0. Specifically, we used *cellranger mkfastq* to convert the raw base calls from the Illumina NovaSeq6000 into FASTQ files. Then we used *cellranger count* to map the reads to the customized combination reference genome of *Macaca mulatta* (Mmul10) and the SIV genome divided into 5 regions (based on NCBI reference sequences M22262.1), along with filtering, barcode counting, and unique molecular identifier (UMI) counting (142). As a result, a gene-cell matrix is generated for each sample. DoubletFinder was used to remove doublets on each of the matrices (143).

For in-depth analysis, we used Partek (St. Louis. MO) Flow version 10.0.22. For scRNA-seq of microglia, we used single-cell QA/QC component to filter out cells with the following attributes: total UMI <400 or >20,000, gene count per cell <300 or >5000, and mitochondrial UMI proportion > 15%. Lymphocytes were removed (based on canonical T-cell and B-cell gene markers), and an average of 5723 cells per animal was identified for analysis. The remaining genes were then filtered by a rhesus monkey to human mapping list (142). Non-protein coding RNAs, mitochondria-encoded genes, ribosomal protein encoding genes, and any gene that had a maximum expression value ý1 was removed, resulting in 12,308 genes for analysis.

For snRNA-seq we used single-cell QA/QC component to filter out cells with the following attributes: total UMI <1000 or >100,000, gene count per cell <700 or >10,000, and mitochondrial UMI proportion > 1%. An average of 5616 nuclei from frontal lobe white matter and 5906 nuclei from the caudate were identified for analysis from each animal. The remaining genes were then filtered by a rhesus monkey to human mapping list (142), and any gene with a maximum expression value ý1 was removed, resulting in 14,143 and 14,379 genes for analysis for the frontal lobe white matter and caudate nucleus, respectively.

Normalization was performed using scTransform (144), followed by principal component analysis (PCA) and then Harmony to eliminate batch effects (145). UMAP (146) and graph-based clustering were then performed, the latter using Louvain clustering (147) with a resolution of 0.3. ANOVA was used for differential expression analysis based on the expression data normalized by log_2_(counts per million plus 1). DEG lists were defined as genes with a false discovery rate of <0.01, a fold change >|1.5|and a minimal expression (least square mean) of at least 50 in one of the conditions or clusters.

GSEA (version 4.1.0) was performed on gene sets for DAM, ARM (both sets taken from Supplemental Table S1 in (148)), and neurodegeneration-related brain myeloid cells (from Supplemental Data S4 in (78)), converting the genes to rhesus gene symbols and eliminating any ribosomal protein genes. To perform GSEA we then used the pre-ranked gene lists from each of the clusters for those genes with an FDR <0.01 on differences between morphine and saline for each cluster, with 1000 permutations to calculate the p-value (149). IPA (150) was performed to generate canonical pathways based on the DEGs between morphine and saline for each cluster.

### Data availability

The snRNA-seq data have been deposited in NCBI GEO, accession # GSE209606. The scRNA-seq data for animals 73T, 75T, 77T, 78T, and 80T have also been deposited in accession # GSE209606. The scRNA-seq data for 74T, 82T, and 83T were previously deposited, accession # GSE195574.

### Statistics

Statistical analyses for studies other than those described in the bioinformatic section above were performed using Prism (version 9.4.0, GraphPad Software, San Diego, CA). Specific tests are indicated in each figure legend.

### Study approval

No new animals were used in these studies, experiments were carried out on archived material from a prior study (57). In that study, macaques were housed in compliance with the Animal Welfare Act and the Guide for the Care and Use of Laboratory Animals in the nonhuman primate facilities of the Department of Comparative Medicine, UNMC, which has been accredited by the AAALAC international. The UNMC Institutional Animal Care and Use Committee reviewed and approved that study under protocol 16-073-FC.

## Author contributions

Conceptualization: HSF, SNB, SB; Investigation: HSF, BM, SGL, KME, PP, SC, AA, GK, SRD; Data – curation: HSF, BM, SRD; Data – formal analysis: HSF, MN, BD, CVF; Project administration: HSF, BM, BGL, JE, CVF; Funding acquisition: HSF, SNB, SB, CVF; Writing – original draft: HSF; Writing – reviewing, editing and revision: HSF, MN, BM, MGL, KME, PP, SC, AA, GK, JE, CG, SRD, CVF, SNB, SB.

## Acknowledgements

This work was supported by grant #s R01DA043164 (to HSF, SB, and SNB) and U01DA053624 (to HSF and SB) from the National Institute of Drug Abuse, P30MH062261 (to HSF and SB) from the National Institute of Mental Health, and R01AI124965 (to CVF) from the National Institute of Allergy and Infectious Diseases, all from the National Institutes of Health. The UNMC DNA Sequencing Core receives support from NIH grants P20GM103427 and P30CA036727. The UNMC Flow Cytometry Research Facility receives state funds from the Nebraska Research Initiative and the NIH grant P30CA036727. The UNMC Bioinformatics and Systems Biology Core is partly supported by NIH awards P20GM103427, P30CA036727, P30MH062261, and U54GM115458. Major instrumentation has been provided by the UNMC Office of the Vice Chancellor for Research, The University of Nebraska Foundation, the Nebraska Banker’s Fund, and by the NIH Shared Instrument Program. We thank Kabita Pandey, Lindsey A. Knight, Alexandra Sheldon, Chase Ochs and Natasha Ferguson for experimental assistance.

---

**Supplemental Figure 1.**
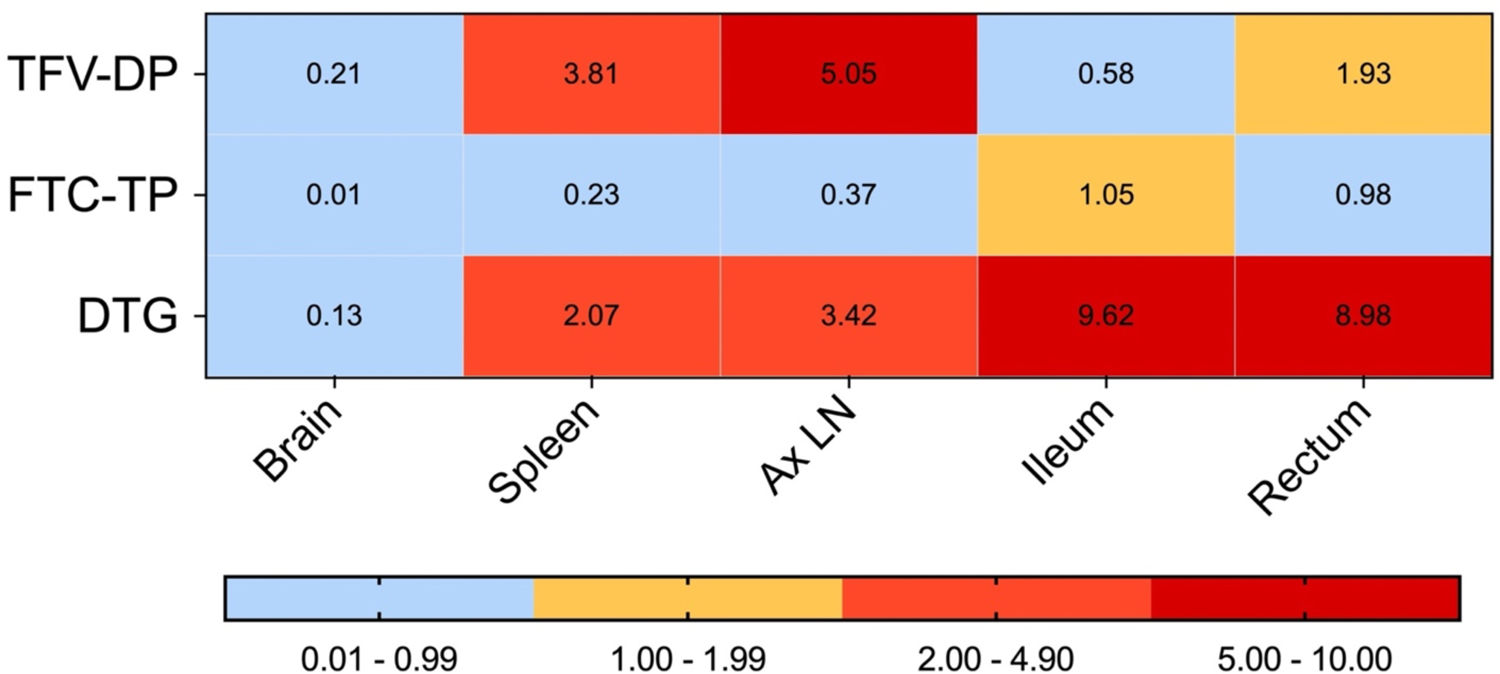
Ratio of tissue concentration of active ARV moiety to the IC_50_ (TFV-DP and FTC-TP) and IC_90_ (DTG). The concentrations in the 5 regions of the brain were averaged for the brain value.

**Supplemental Figure 2.**
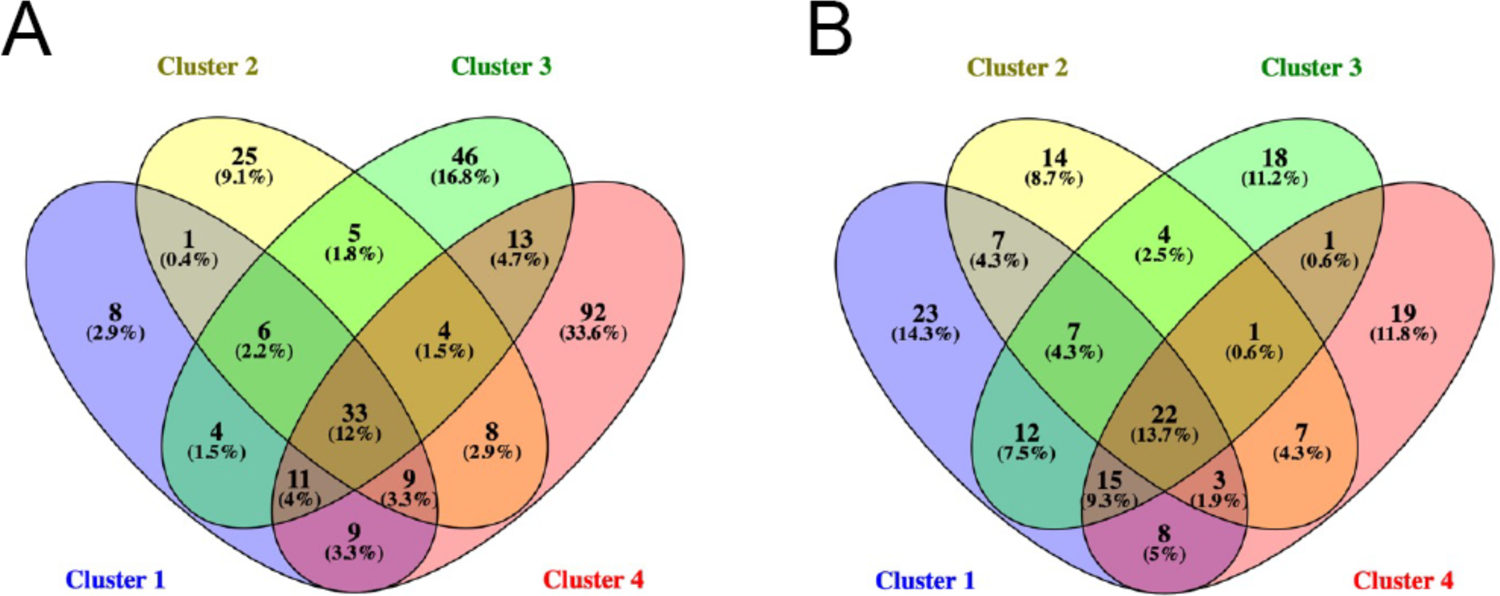
Common genes are differentially regulated by morphine in microglia clusters. Venn diagram showing the intersections of the **A)** up-regulated and **B)** down-regulated (right) DEGs found in microglia clusters 1-4.

## Supplemental Table Legends

**Supplemental Table 1. Plasma biomarkers.** Values before and after morphine administration for the morphine group, and before and 2, 5, 22, and 35-weeks after SIV inoculation are given for all monkeys. For statistical purposes values <LLOQ were one-half of the LLOQ.

**Supplemental Table 2. DEGs between the clusters.** FDR q-value, fold-change, and least-square mean (LSMean) values for each cluster compared to the other 5 clusters. Each cluster is shown on a separate worksheet, DEGs lists were filtered for an FDR <0.01, fold change >|1.5|and a minimal LSMean expression of at least 50 in either the indicated cluster or the combination of the other clusters.

**Supplemental Table 3. DEGs within each cluster comparing the morphine group to the saline group.** The rhesus gene IDs and the human homologues are indicated, along with the FDR q-value, fold-change, and least-square mean (LSMean) values for the morphine group and the saline group within each cluster. Each cluster is shown on its indicated worksheet, DEGs lists were filtered for an FDR <0.01, fold change >|1.5|and a minimal LSMean expression of > 50 in either the morphine or saline group within the cluster.

**Supplemental Table 4. Gene sets used for GSEA.** The published lists of human genes for the gene sets are listed, followed by the mapping to rhesus homologues. The gene names highlighted in grey code for ribosomal proteins that were not considered in the GSEA.

**Supplemental Table 5. IPA analysis.** The canonical pathways assessed for the DEGs found due to morphine treatment are shown on a separate worksheet for each cluster. Highlighted are those with a p-value <0.05 and a Z-score> 1.

